# Transcriptional Network Analysis on Brains Reveals a Potential Regulatory Role of *PPP1R3F* in Autism Spectrum Disorders

**DOI:** 10.1101/330449

**Authors:** Abolfazl Doostparast Torshizi, Jubao Duan, Kai Wang

## Abstract

**Objective:** This study aims at identifying master regulators of transcriptional networks in autism spectrum disorders (ASDs).

**Results:** With two sets of independent RNA-Seq data generated on cerebellum from patients with ASDs and control subjects (N=39 and 45 for set 1, N=24 and 38 for set 2, respectively), we carried out a network deconvolution of transcriptomic data, followed by virtual protein activity analysis. We identified PPP1R3F (Protein Phosphatase 1 Regulatory Subunit 3F) as a master regulator affecting a large body of downstream genes that are associated with the disease phenotype. Pathway enrichment analysis on the identified targets of *PPP1R3F* in both datasets indicated alteration of endocytosis pathway. This exploratory analysis is limited by sample size, but it illustrates a successful application of network deconvolution approaches in the analysis of brain gene expression data and generates a hypotheses that may be further validated by large-scale studies in the future.

## Introduction

Autism Spectrum Disorders (ASD) comprise a set of highly inheritable neurodevelopmental conditions characterized by impairments in social communication, repetitive behaviors and restricted interests [1, 2]. ASDs are estimated to affect 1 in 68 children in the United States, and boys are 4.5 times more likely than girls to develop ASDs [3]. Several studies showed that the heritability of autistic phenotypes is estimated to be around 90% [4, 5]. The number of genes potentially implicated in ASDs is rapidly growing, mainly from large-scale genetic studies such as next generation sequencing (NGS) [6-10] and genome wide association studies (GWAS) [11-13]. Although these genetic studies have substantially advanced our understanding of the etiology of ASDs, the underlying molecular mechanisms remain elusive [14]. Transcriptome analysis is gaining momentum as a complementary approach to genetic association studies [14], and can help us understand the molecular pathophysiology of ASDs in a more systematic and mechanistic manner.

A number of studies have been conducted to evaluate whole-genome gene expression that may contribute to the onset of ASD. In a large-scale RNA-Seq effort, matched brain regions from subjects affected with ASDs and controls were utilized to identify neuronal genes which are strongly dysregulated in cortical regions [14]. It was noted that a module of expressed genes in microglia was negatively correlated with a module of differentially expressed neuronal genes, implicating correspondence of dysregulated microglial responses with activity-dependent genes in autism brains [14]. Utilizing microarray technology, Voineagu et al. [15] demonstrated consistent differences in transcriptome organization between autistic/normal human brain tissues using gene co-expression network analysis. In their study, they report consistent differences in regional patterns of gene expression in several brain regions such as frontal cortex, suggesting abnormalities in cortical patterning [15]. However, besides extracting co-expressed gene modules, they did not investigate potential molecular drivers of such expression patterns. Despite applications of co-expression network approaches in the inference of regulatory machinery in ASD [16], state-of-the-art network approaches such as information-theory based methods and network deconvolution methods are barely adopted in this area. Network deconvolution methods have shown great merits in a wide range of disease or mechanistic studies such as prostate differentiation [17] and cancers [18]. These methods can overcome limitations of other network-based methods such as: connecting genes having indirect interactions leaving their mutual causal effects aside, overfitting when dealing with small number of samples, suffering from the curse of dimensionality, and not being able to reverse-engineer the mammalian genome-wide cellular networks [19]. Using available gene expression data, these methods can provide deep biological insights into the underlying transcription circuitry of diseases and illustrate molecular connections and potential regulation drivers at a systems level. As an example, with transcriptional network deconvolution approach, we have recently provided novel insights on Post-Traumatic Stress Disorder (PTSD) [20] by identifying several genes as drivers of innate immune function. In the current study, we used ARACNe (Algorithm for Reconstruction of Accurate Cellular Networks) [21] as a versatile tool to deconvolute cellular networks. In this approach, first gene-gene co-regulatory patterns are identified using the information theoretic measure of mutual information. Next, the constructed networks are pruned by removing indirect connections where two genes are co-regulated through one or more intermediaries. This allows us to keep connections bearing significantly high probabilities of representing direct interactions or mediated interactions through post-transcriptional regulation. The constructed networks were then used to infer the activity degrees of the hub genes within the network.

In this study, using two of the largest transcriptomic datasets of postmortem brain tissues from ASD individuals and control subjects by Parikshak *et al* [16] and Gupta *et al* [14], we reconstructed the transcriptional networks followed by virtual protein activity analysis, to identify “master regulators” (MRs) that may differentially regulate the expression levels of multiple downstream genes in the cerebellum region of ASD patients and controls.

## Main Text

## Methods

In this study, after constructing the transcriptional networks, we have used a probabilistic algorithm called VIPER (Virtual Inference of Protein-activity by Enriched Regulon analysis [18]). VIPER aims at inferring the protein activity of a MR by a systematic analysis of the expression patterns of its targets (regulons). VIPER directly integrates target mode of regulation indicating whether targets are repressed or activated given the statistical confidence in regulator-target interactions and target overlap between different regulators in order to obtain the enrichment of a protein regulon in differentially expressed genes [20]. VIPER has advantages over the existing approaches such as T-profiler [22], gene set enrichment analysis (GSEA) [23], and Fisher‘s exact test [24] in that it uses a fully probabilistic enrichment analysis framework that supports seamless integration of genes with different likelihoods of representing activated, repressed or undetermined targets.

Both datasets contain multiple brain regions including cerebellum, which is relevant for ASDs since specific cerebellar zones can affect neocortical substrates for social interaction and cognitive functions such as language and executive functions [25-27]. We reasoned that in the same brain region, there should be highly active proteins whose expression regulate a large set of target genes and such patterns should be replicated in an independent RNA-Seq data. Our preliminary finding indicates *PPP1R3F* (Protein Phosphatase 1 Regulatory Subunit 3F) as a potential master regulator (MR) to exert large-scale regulatory effects on a body of genes in ASD. The general framework of the in-silico experiments is illustrated in Figure 1. We further explored the relevance of this gene to ASD and generated a testable hypothesis on how the dysregulation of this gene can influence ASD pathogenesis.

**Figure 1.**
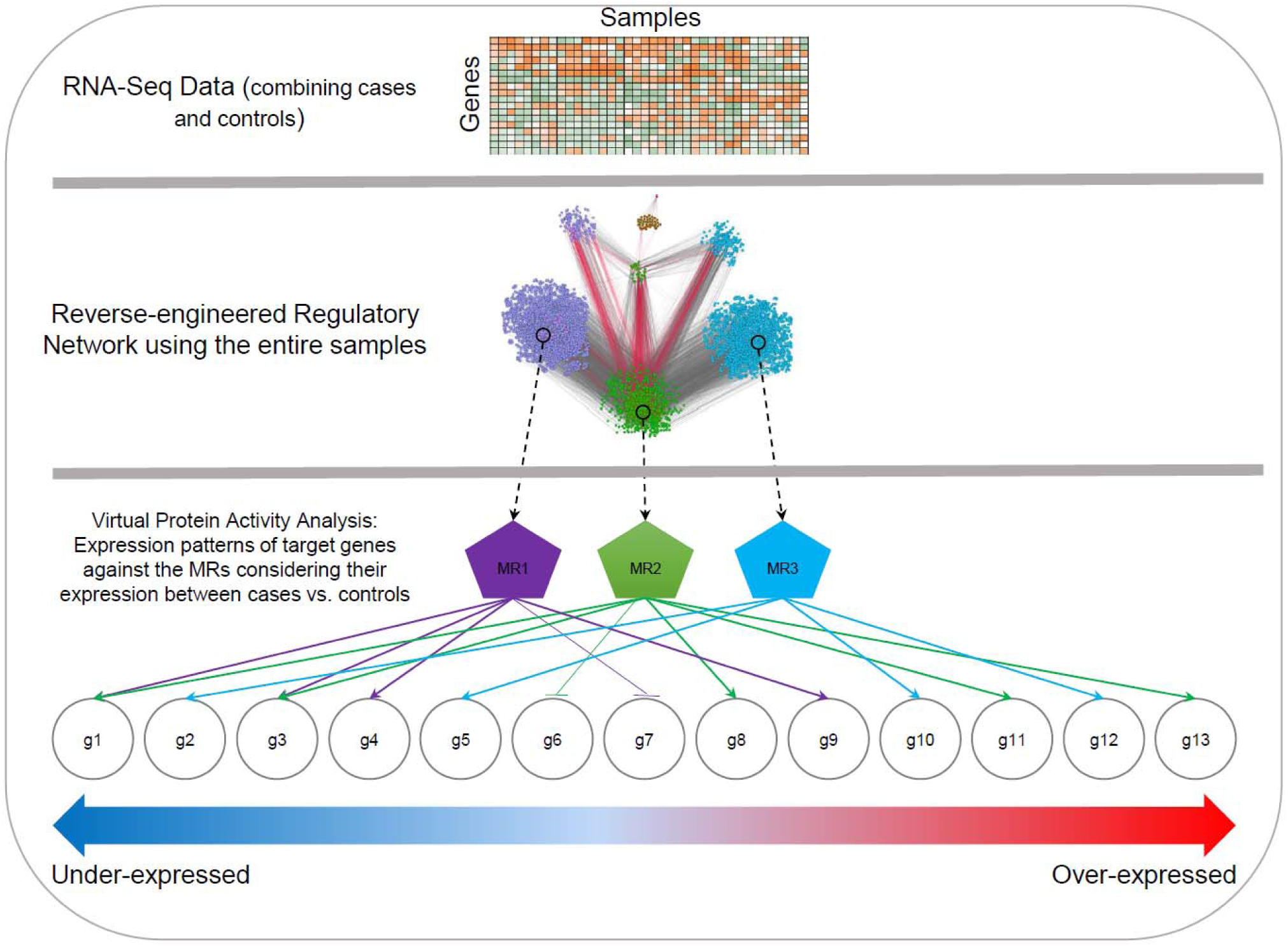
The overall process of network construction and virtual protein activity analysis to identify a master regulator

## Results and discussion

We first used the data from Parikshak *et al* [16] to construct the regulatory networks. This data is part of a large RNA-Seq repository on post-mortem human brain tissue (39 cases versus 45 controls) from cerebellum, frontal cortex, temporal cortex, prefrontal cortex, and visual cortex. During the process of network deconvolution (see Methods in Additional File 1), pairwise Mutual Information (MI) between all of the available transcripts were obtained. Next, the constructed network was trimmed to remove genetic intermediaries, resulting in potential direct connections between MRs and their targets (we used the recommended p-value threshold of 1e-8, as a measure of confidence of regulatory relationships between two genes [21]). This analysis yielded a repertoire of 672,973 interactions, 23,935 regulators, and 24,847 targets in the constructed network using the dataset from Parikshak *et al* [16]. We similarly analyzed the second dataset from Gupta *et al* [14], a RNA-Seq data of post-mortem brain tissues where cerebellum samples were much more than samples from other brain regions. Using the same network construction settings on this dataset [14] containing 24 cases and 38 controls, we deconvoluted a network of 297,870 interactions containing 12,040 regulators and 12,529 targets. The constructed networks from both datasets were provided in Additional Files 3 and 4, respectively.

After applying VIPER on the constructed networks using both datasets, we compared the list of highly significant MRs at FDR ≤ 0.05. We identified *PPP1R3F* (Protein Phosphatase 1 Regulatory Subunit 3F) as the only shared MR between the two data sets; given the small sample size, it is possible that our analysis was under-powered and may have missed other relevant MRs in ASDs. Figure 2 illustrated how downregulation of this MR influences the expression of its regulons in both constructed networks.

**Figure 2.**
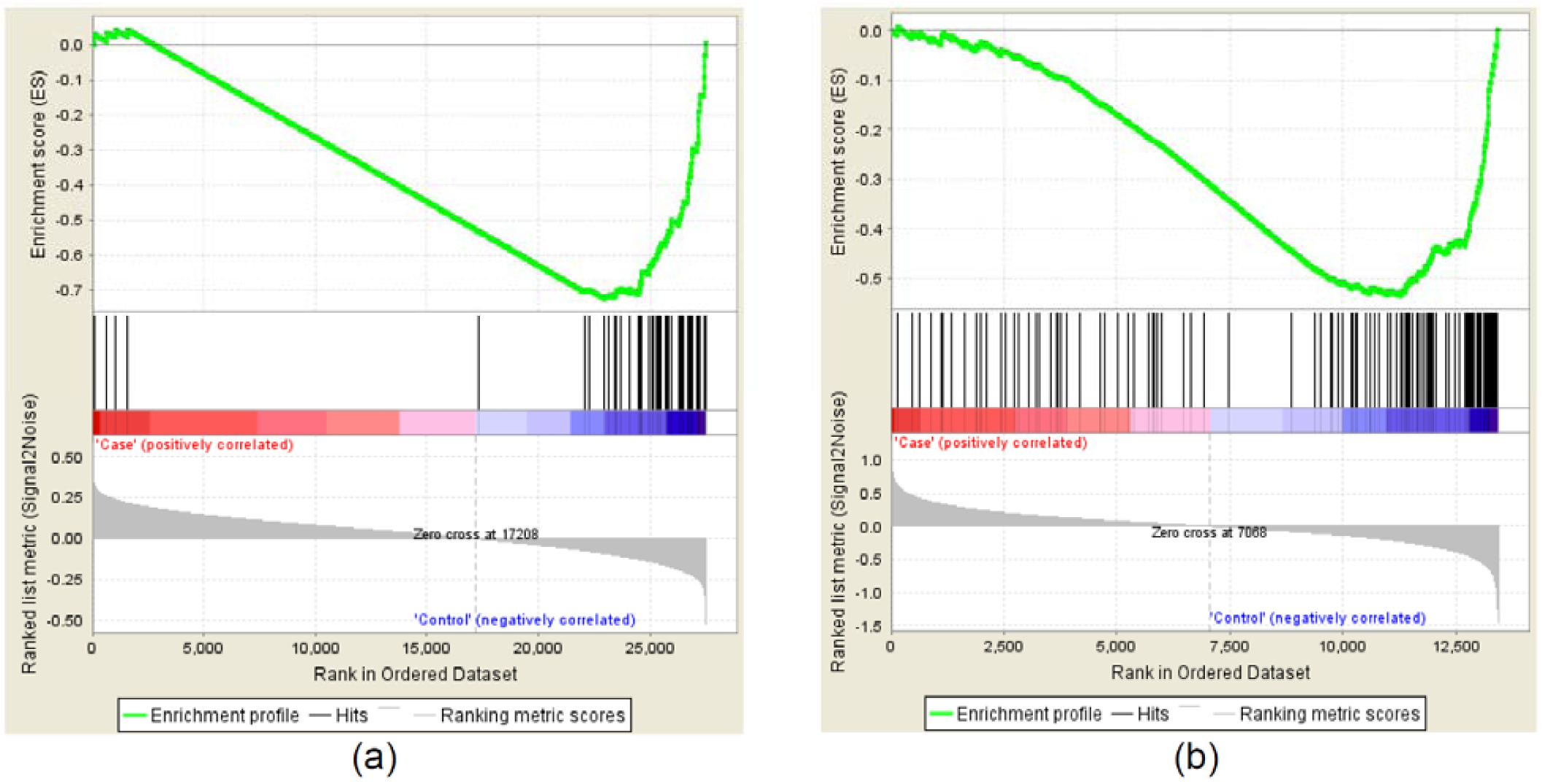
Gene set enrichment analysis (GSEA) of *PPP1R3F* targets in the constructed networks using the data by (a) Parikshak *et al* [16] and (b) Gupta *et al* [14]. Black bars in the both figures depict the rank of the *PPP1R3F* targets in terms of correlation with the phenotype among the entire list of genes in the both datasets.

*PPP1R3F* is one of the type-1 protein phosphatase (PP1) regulatory subunits which is found to be important in neuronal activities and has been implicated in carrying rare mutations in autistic individuals [28]. A systematic resequencing of X-chromosome synaptic genes in a group of individuals with ASD (122 males and 20 females) has identified a rare non-synonymous variant in *PPP1R3F* that can predispose to developing ASDs [28]. This potentially damaging variant, c.733T> C, was observed in a boy with a diagnosis of Asperger Syndrome and was transmitted from a mother who suffered from learning disabilities and seizures [28]. Protein phosphorylation is a key mechanism by which cells regulate transduction pathways and PPP1 family enzymes are associated with dephosphorylation of several proteins such as TGF-b cascade [29].

To further probe the biological relevance of the predicted *PPP1R3F* network to ASD, we examined the overlaps between *PPP1R3F* regulons and known candidate genes implicated in ASD and its related disorders (Table 1). The most significant overlap was found with SFARI gene list [30] (P=0.0008), followed by overlap with an intellectual disability database gene list (P=0.072) [31]. The overlaps with other ASD candidate gene lists also showed trends towards to being significant). These results suggest the potential relevance of the predicted *PPP1R3F* network to ASD.

**Table 1.**
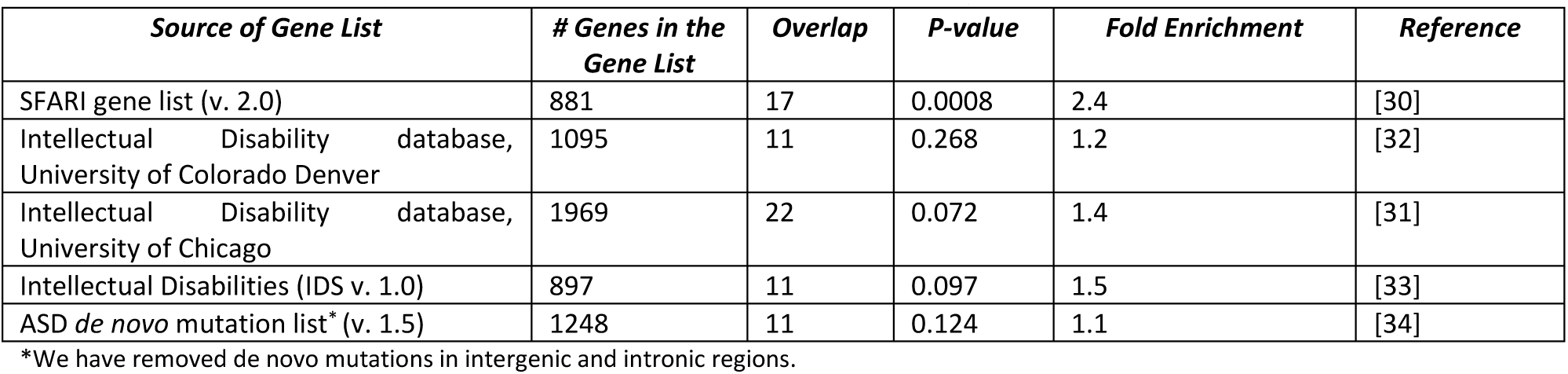
The overlap of the identified *PPP1R3F* regulons from both datasets (n= 177 genes) with several candidate gene lists for ASDs and ID (Intellectual Disability). P-values are calculated by two-sided Fisher‘s exact test.

| Source of Gene List | # Genes in the Gene List | Overlap | P-value | Fold Enrichment | Reference |
| --- | --- | --- | --- | --- | --- |
| SFARI gene list (v. 2.0) | 881 | 17 | 0.0008 | 2.4 | [30] |
| Intellectual Disability database, University of Colorado Denver | 1095 | 11 | 0.268 | 1.2 | [32] |
| Intellectual Disability database, University of Chicago | 1969 | 22 | 0.072 | 1.4 | [31] |
| Intellectual Disabilities (IDS v. 1.0) | 897 | 11 | 0.097 | 1.5 | [33] |
| ASD *de novo* mutation list* (v. 1.5) | 1248 | 11 | 0.124 | 1.1 | [34] |

\*We have removed *de novo* mutations in intergenic and intronic regions.

*PPP1R3F* is a sex-linked gene, so we attempted to account for the differences between the expression of *PPP1R3F* in male and female samples with ASDs. In the Parikshak *et al* data set (from [16]) there were 32 males and 7 females with ASDs while there were 39 male controls compared to 6 female controls. The gender information is not available on the Gupta *et al* dataset [14]. We found no difference of *PPP1R3F* expression between male and female samples in the Parikshak *et al* dataset [16] (FDR=0.644; two-side *t*-test), but this may be due to small sample size. Nevertheless, to account for possible role of sex-related gene expression in the structure of the constructed network, we re-constructed the regulatory network using only male samples in the Parikshak *et al* dataset [16] (that is, 32 cases and 39 controls). Following the virtual protein activity analysis, we observed that *PPP1R3F* remained as a significant MR (VIPER enrichment P-value=0.0186). This finding indicated that *PPP1R3F* acts independently from potential sex-based gene expression differences, and thus our finding of *PPP1R3F* as a MR was not an artifact of sex biases. Additionally, we conducted the same analytical experiments on the gene expression data from pre-frontal cortex. *PPP1R3F* was not identified as a significant MR in this brain region (activity FDR=0.1364). We should note that the number of samples from other brain regions were too small to be used for network analysis. Our finding suggests the potential role of *PPP1R3F* in developing ASDs upon regulatory control of a large body of genes in the cerebellum region of the brain.

We next conducted pathways enrichment analysis on the *PPP1R3F* regulons from both constructed networks separately, and observed the targets to be enriched for endocytosis pathway in the Parikshak *et al* dataset [16] (FDR=0.005, fold enrichment= 8.26) and the Gupta *et al* dataset [14] (FDR=0.0008, fold enrichment= 8.42). “Endocytosis” is the only significantly enriched pathway on both data sets. Combining both sets of targets totaling 177 genes (Supplementary Figure 1 in Additional File 2 and Additional File 5), the enrichment of endocytosis pathway was even more significant (FDR=4.85e-04, fold enrichment= 8.97).

Since ASDs are commonly recognized as brain disorders, we further examined whether the identified MR is mainly expressed in the brain. We looked up *PPP1R3F* in GTEx consortium portal [35], and found that compared to other tissues, *PPP1R3F* is predominantly expressed in various brain regions such as frontal cortex and cerebellum (Supplementary Figure 2 in Additional File 2). We checked BrainSpan Atlas of the Developing Human Brain (http://brainspan.org) to see whether *PPP1R3F* is highly expressed in prenatal or postnatal stages. *PPP1R3F* was not expressed until 37 weeks post-conception. While remaining unexpressed in some brain regions, it is modestly expressed in four-month postnatal stage in particular brain regions. We further probed the expression of each of the 177 targets of *PPP1R3F* in GTEx and identified the tissues in which they are highly expressed. 89 genes out of 177 were highly expressed in various regions of the brain, compared to other tissues (*P*-value from Fisher‘s exact test= 5.51 × 10^−5^, number of protein coding genes in GTEx = 20900, number of protein coding genes highly expressed in the brain in GTEx=7528). The significant overlap of the expressed *PPP1R3F* target genes with the total number of highly expressed genes in the brain partially supports the pathophysiological relevance of *PPP1R3F* to ASDs.

## Conclusions

In this study, we performed exploratory analysis on two small-scale RNA-Seq data sets, and used a network deconvolution algorithm to reverse engineer regulatory networks. By further applying virtual protein activity analysis on both networks, we identified *PPP1R3F* as a MR of a regulon consisting of 177 targets that are differentially expressed between ASD patients and controls. Gene set enrichment analysis on the *PPP1R3F* regulons suggested that *PPP1R3F* may exert its functional effects through regulating the endocytosis, a pathway that has been previously implicated in neuropsychiatric disorders [36].

## Limitations

We acknowledge that our study is limited by the small sample size (due to the scarcity of brain tissues), and the results thus need further replications. Nonetheless, our study generates a testable hypothesis that may be validated by large-scale studies in the future. Additionally, further experimental validation of the regulatory effects of *PPP1R3F* on its downstream targets as predicted by our network analysis may provide novel insights on possible pathophysiological role of *PPP1R3F* as a MR of ASD gene network.

## Additional files

Additional file 1. Detailed explanation of the methods being used in this study.

Additional file 2. Supplementary figures.

Additional file 3. The constructed networks from the Parikshak *et al* dataset [16].

Additional file 4. The constructed networks from the Gupta *et al* dataset [14].

Additional file 5. The list of the combined set of target genes of *PPP1R3F.*

## Declarations

### Ethics approval and consent to participate

Not Applicable

### Consent for publication

Not Applicable

### Availability of data and materials

All data used in this study are available from references [14, 16].

### Competing Interests

The authors declare no conflict of interests.

### Funding

This study is supported by NIH grant MH108728 to K.W.

#### Acknowledgment

The authors would like to thank the PsychENCODE consortium for providing the data. Data were generated as part of the PsychENCODE Consortium, supported by: U01MH103339, U01MH103365, U01MH103392, U01MH103340, U01MH103346, R01MH105472, R01MH094714, R01MH105898, R21MH102791, R21MH105881, R21MH103877, and P50MH106934 awarded to: Schahram Akbarian (Icahn School of Medicine at Mount Sinai), Gregory Crawford (Duke), Stella Dracheva (Icahn School of Medicine at Mount Sinai), Peggy Farnham (USC), Mark Gerstein (Yale), Daniel Geschwind (UCLA), Thomas M. Hyde (LIBD), Andrew Jaffe (LIBD), James A. Knowles (USC), Chunyu Liu (UIC), Dalila Pinto (Icahn School of Medicine at Mount Sinai), Nenad Sestan (Yale), Pamela Sklar (Icahn School of Medicine at Mount Sinai), Matthew State (UCSF), Patrick Sullivan (UNC), Flora Vaccarino (Yale), Sherman Weissman (Yale), Kevin White (UChicago) and Peter Zandi (JHU). We would also like to thank the providers of the Gupta *et al* dataset (The Arking Lab at the McKusick-Nathans Institute of Genetic Medicine of Johns Hopkins University) to generate the brain gene expression data and make the data freely available.

## Author‘s Contributions

A.D.T. designed the experiments, conducted the analysis and computations, and wrote the manuscript. J.D. advised on data analysis and edited the manuscript. K.W. supervised the research, and edited the manuscript.

